# The effect of semantic brightness on pupil size: A replication with Dutch words

**DOI:** 10.1101/689265

**Authors:** Sebastiaan Mathôt, Leevke Sundermann, Hedderik van Rijn

## Abstract

Theories of embodied language hold that word processing is automatically accompanied by sensory and motor simulations. For example, when you read the word ‘sun’, a sensory simulation of brightness as well as a motor simulation of pupil constriction would be automatically triggered. Consistent with this notion, Mathôt, Grainger, and Strijkers (2017) found that the eye’s pupil was slightly smaller after reading single words that were associated with brightness (e.g. ‘sun’) as compared to darkness (e.g. ‘night’); that is, the pupil light response was modulated by the semantic brightness of words. However, (other) key findings within the field of embodied language have proven difficult to replicate, and we therefore felt that it was crucial to replicate the effect of semantic brightness on pupil size. To this end, we conducted a close-but-non-identical replication of two key experiments from Mathôt, Grainger, and Strijkers (2017): one experiment with visually presented words, and one experiment with spoken words. Both experiments were successfully replicated. We propose that cognitive modulations of the pupil light response reflect activity in visual brain areas; therefore, the effect of semantic brightness on pupil size can be used as a marker for the involvement of visual brain areas in language processing, and thus to address a wide variety of key questions within psycholinguistics.

## The Effect of Semantic Brightness on Pupil Size

The pupil light response (PLR) is a constriction of the eye’s pupil in response to brightness, and a dilation of the pupil in response to darkness (reviewed in Mathôt, 2018). The PLR is thought to be a mechanism that optimizes vision based on the amount of light that is available (Campbell & Gregory, 1960; M. Lombardo & Lombardo, 2010; Watson, 2013). Specifically, small pupils have the advantage of suffering less from optical distortions that reduce visual acuity; therefore, small pupils facilitate discrimination of fine detail (acuity). However, small pupils have the disadvantage of reducing the amount of light that reaches the retina; therefore, small pupils impair detection of faint stimuli (sensitivity). One of the functions of the PLR is likely to seek an optimal point in the trade-off between visual acuity and sensitivity by reducing pupil size when sufficient light is available, and increasing pupil size when there is not. Traditionally, this was considered a low-level physiological reflex that is not affected by higher-level cognition.

However, evidence for cognitive modulations of the PLR dates back many decades (Bárány & Halldén, 1948; Harms, 1937; Lowe & Ogle, 1966), and has recently attracted renewed attention (reviewed in e.g. Binda & Murray, 2014; Mathôt & Van der Stigchel, 2015). These classic studies used binocular rivalry, a technique in which a different stimulus is shown to each eye. This results in visual awareness alternating between the two eyes: Sometimes the participant is aware of what is presented to the right eye, and sometimes of what is presented to the left eye. The crucial finding from these studies is that the PLR is strongest to stimuli that are presented to the eye that currently dominates awareness: a clear demonstration that the PLR is modulated by cognition (for similar recent studies, see Fahle, Stemmler, & Spang, 2011; Naber, Frassle, & Einhauser, 2011; Sperandio, Bond, & Binda, 2018).

In recent years, studies have shown that the PLR is modulated by a wide range of cognitive factors: eye-movement preparation, such that a pupil constriction is prepared along with an eye movement towards a bright object (Ebitz, Pearson, & Platt, 2014; Mathôt, van der Linden, Grainger, & Vitu, 2015); covert visual attention, such that the PLR is strongest to stimuli presented within the focus of attention (Binda, Pereverzeva, & Murray, 2013a; Mathôt, van der Linden, Grainger, & Vitu, 2013; Naber, Alvarez, & Nakayama, 2013); visual working memory (vWM), such that the pupil is smaller when a bright, as compared to a dark, stimulus is maintained in vWM (Husta, Dalmaijer, Belopolsky, & Mathot, 2018); subjective interpretation, such that the pupil is smaller when people interpret a picture as depicting something bright, such as a sun, regardless of the actual brightness of the picture (Binda, Pereverzeva, & Murray, 2013b; Laeng & Endestad, 2012; Naber & Nakayama, 2013); and mental imagery, such that the pupil constricts when people merely imagine a bright object or scene (Laeng & Sulutvedt, 2014). Taken together, these studies show clearly that the PLR, while not being under direct voluntary control, is modulated by higher-level cognition.

Building on these studies, Mathôt, Grainger, & Strijkers (2017) recently used the PLR to test one of the central claims of theories of embodied language, which is that word comprehension is accompanied by sensory and motor simulations that are associated with the word’s referent (Glenberg & Gallese, 2012; Pulvermüller, 2013). For example, reading the word ‘sun’ would be accompanied by a sensory simulation of a bright ball of fire in the sky, as well as a motor simulation of pupil constriction. According to theories of embodied language, such sensory and motor simulations are an important way, if not the only way, in which we process and understand language (but for an opposing view, see Mahon & Caramazza, 2008). Mathôt et al. (2017) hypothesized that these simulations should result in a slight-yet-measurable constriction of the pupil when participants read or heard words that were associated with brightness (e.g. ‘sun’ or ‘light’), as compared to words that words that were associated with darkness (e.g. ‘night’ or ‘shadow’). The results from two experiments confirmed this: Pupil size was indeed modulated by the semantic brightness of words.

We feel that it is important to replicate the results from Mathôt et al. (2017) for two main reasons. First, the original study used a limited set of French words, and we therefore want to establish whether the results generalize to a different set of words, in this case even a different language (Dutch). (But still limited in number, because it is difficult to find many brightness- and darkness-related words that are matched on crucial visual and lexical properties.) Second, Papesh (2015) recently reported a failure to replicate one of the landmark behavioral studies in the field of embodied language (Glenberg & Kaschak, 2002). Therefore, it is important to verify the reliability of studies within this field, most of which have not yet been the focus of (published) replication attempts. Yet some of these studies have greatly influenced the theoretical debate (e.g. Glenberg & Kaschak, 2002); and other, more recent studies may do so in the future (e.g. Mathôt et al., 2017).

We conducted two experiments that were close replications of the auditory and visual experiments from Mathôt et al. (2017). The main difference between the current study and the original study is the language: We used Dutch, rather than French, words. To facilitate direct comparison, we conducted similar (but not identical) analyses, and visualized the results in the same way. In addition, we also collected concreteness ratings for all words; this is important because concrete words require less cognitive processing than abstract words (Schwanenflugel, 2013), and concreteness could therefore have been in a confound in Mathôt et al. (2017). To foresee the result, we successfully replicated the effect of semantic brightness on pupil size with both visual and auditory stimuli, and also after controlling for concreteness.

## Methods

The paradigm was almost identical to that used in Mathôt et al. (2017), with the notable difference that we used Dutch words, rather than French words. The experiments were approved by the local ethics committee of the University of Groningen (IRB codes 16129-S-N, 17228-S-N, 17229-S-N).

### Stimulus selection

We manually selected 24 brightness-related words, 24 darkness-related words, 24 control words (i.e. not clearly related to brightness or darkness), and 16 animal names from the Dutch lexicon project (Keuleers, Diependaele, & Brysbaert, 2010). The brightness- and darkness-related words were carefully matched on number of letters (bright: *M* = 5.96; dark: *M* = 5.83; range: 3 - 11) and lexical frequency (bright: *M* = 2,839 occurrences per million; dark: *M* = 2,781; range = 52 - 25,976). For the Visual experiment, we rendered each word as an image, using a monospace font. Next, we resized the images to match the summed pixel intensity of the brightness- and darkness-related words (bright: *M* = 1.17 × 10^6^ arbitrary units; dark: *M* = 1.17 × 10^6^). For the Auditory experiment, we rendered each word as a sound file, using the Mac OS text-to-speech synthesizer (`say`). The control words and animal names were not matched; therefore, we consider only the brightness- and darkness-related words in the statistical analyses described below.

### Participants

31 participants took part in the Visual experiment (one more than planned due to excess sign-ups); 30 other participants took part in the Auditory experiment. Our sample size was based on Mathôt et al. (2017). Participants were compensated with course credit, had normal or corrected-to-normal vision, and were students from the University of Groningen. All but one of the participants were native speakers of Dutch. For privacy, we did not collect specific demographic information.

### Software and apparatus

Eye movements and pupil size were recorded with an EyeLink 1000 (SR Research, Missisauga, Ontario, Canada), a video-based eye tracker with a temporal resolution of 1000 Hz. Testing took place in a dimly lit room. The experiments were implemented with OpenSesame (Mathôt, Schreij, & Theeuwes, 2012), using the Expyriment backend (Krause & Lindemann, 2014) and PyGaze for connecting to the eye tracker (Dalmaijer, Mathôt, & Van der Stigchel, 2014).

### Procedure

Each trial started with a dark-gray central fixation dot, presented against a gray background, for 3 s. Next, the target word was presented. In the Visual experiment, the word was presented visually in a dark-gray font (slightly darker than the background) for 3 s or until the participant pressed a key. In the Auditory experiment, the word was uttered by a synthesized voice through a set of headphones. Participants were instructed to press the spacebar when they saw or heard an animal name, and to withhold a response otherwise, in which case the experiment continued after 3 s (i.e., semantic categorization). This ensured that participants processed the meaning of the words, without performing any overt action in response to the words of interest that would otherwise affect the pupil response. Finally, after the presentation of the word, a dim red or green fixation dot was shown for 500 ms to give feedback after respectively an incorrect or a correct response.

The experiment started with 10 practice trials, for which we selected an additional 8 non-animal names and 2 animal names, and which are not further analyzed. Next, there were 88 trials, divided over 4 blocks. The order of the words was randomized.

## Results

Our analysis is similar to that of Mathôt et al. (2017), but deviates in several ways that are explained below, to adhere more closely to the more recent guidelines outlined in Mathôt, Fabius, Heusden, & Stigchel (2018). The most notable differences are that, in the current study, we report pupil size in millimeters of diameter (as opposed to arbitrary units), that we use subtractive (as opposed to divisive) baseline correction, and that we use a slightly different window for determining the mean pupil response for the individual participant and item analyses.

### Pre-processing and outlier removal

We analyzed pupil size during the 3 s after word onset, and then performed the following pre-processing in this order: We reconstructed blinks and missing data using cubic-spline interpolation (Mathôt, 2013). We downsampled the signal to 100 Hz. We converted pupil size from arbitrary units to millimeters of diameter; for this we used a conversion formula that we determined by recording the size of artificial pupils (dark circles printed on white paper) with a known diameter. We used mean pupil size during the first 50 ms after word onset as a baseline (i.e., close to the start of the relevant interval, but well below the minimum latency of the pupil response, which is around 220 ms; Ellis, 1981). We visually inspected the distribution of baseline pupil sizes to determine sensible minimum and maximum values, which we found to be 2 mm and 6.5 mm. We then excluded all trials in which baseline pupil size fell outside of this range, or in which baseline pupil size could not be determined due to missing data (Visual experiment: 3.3%; Auditory experiment: 6.1%). Finally, we subtracted baseline pupil size from all samples (i.e., subtractive baseline correction).

### Behavioral results

Response accuracy was 94% in the Visual experiment and 93% in the Auditory experment. The mean correct response time to animal names was 989 ms in the Visual experiment and 1,276 ms in the Auditory experiment. (This was slightly higher than the mean correct response time in Mathôt et al. (2017), which was 790 ms for the Visual experiment and 1,067 ms for the Auditory experiment.)

### Pupil-size results

Figure 1 shows mean pupil size for correct trials over time after word onset, separately for brightness-related words, darkness-related words, and control words. (Control words were not matched to the Darkness- and Brightness-Related words, and therefore triggered slightly different overall pupil responses, especially in the Visual modality. For this reason, control words are not included in the statistical analyses, cf. Mathôt et al. (2017).)

**Figure 1.**
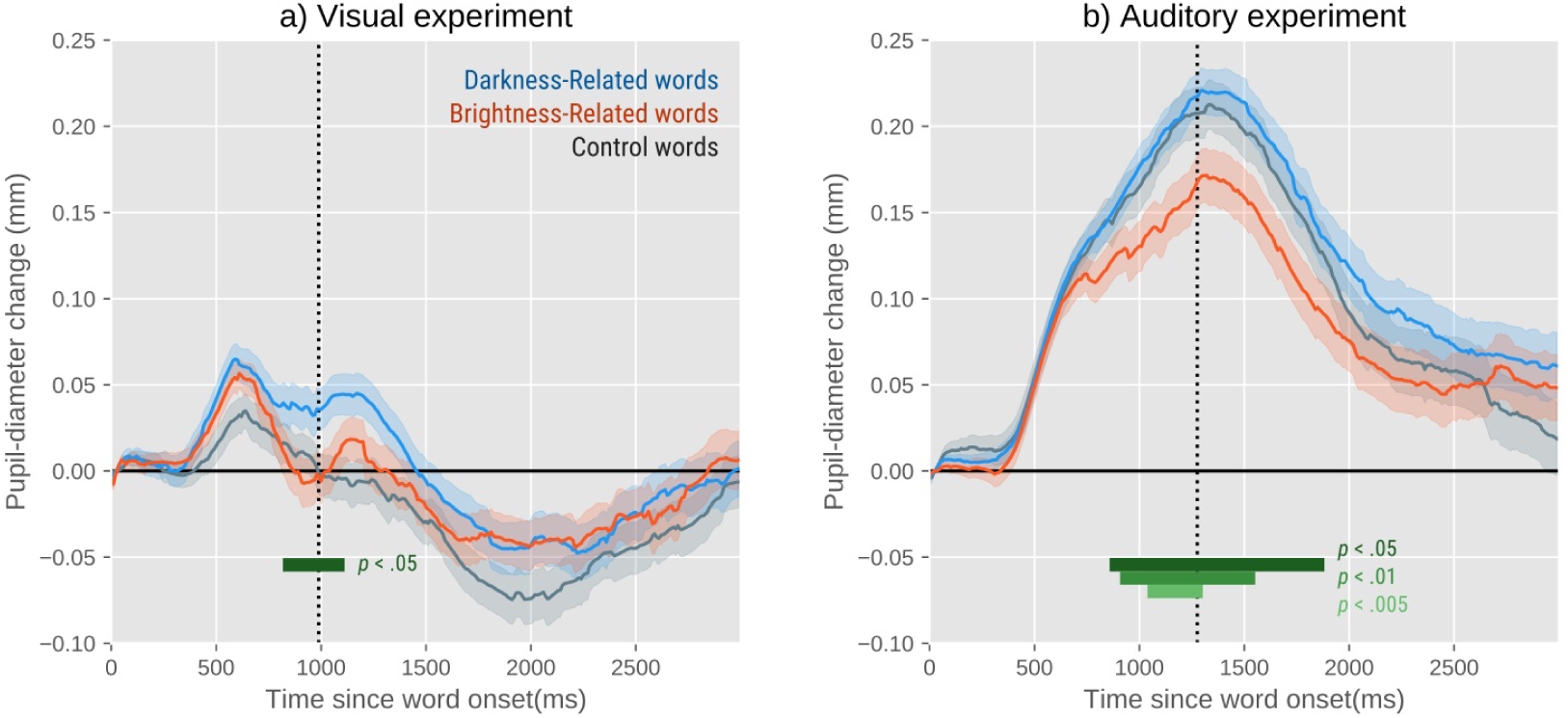
After seeing (a) or hearing (b) a darkness-related word (blue lines), the pupil is slightly larger than after seeing or hearing a brightness-related word (orange lines). The significance values correspond to the contrast between brightness- and darkness-related words.

To test for differences in pupil size, we conducted a linear-mixed effects analysis with Baseline-Corrected Pupil Size as dependent measure, Semantic Brightness (Bright v Dark) as fixed effect, and by-participant random intercepts. We did not include by-participant random slopes, because this regularly caused failures to converge. This analysis was done separately for every 10 ms time bin, with mean pupil size in that bin as dependent measure. We considered an effect to be reliable when *p* < .05 (estimated through *t* > 1.96) for at least 200 contiguous milliseconds (cf. Mathôt et al., 2017). However, we emphasize overall patterns over significance values.

In the Visual modality (Figure 1a), the pupil initially dilates with a peak after about 600 ms; this is an orienting response (Mathôt, Dalmaijer, Grainger, & Van der Stigchel, 2014; Wang & Munoz, 2015), and is not notably modulated by word meaning. Next, there is a second period of dilation with a peak after about 1,200 ms; this presumably reflects higher-level processing (Mathôt, 2018), and is modulated by word meaning (850 - 1080 ms) such that the pupil is larger after darkness-related words than after brightness-related words. Finally, there is a prolonged overall pupil constriction that results from visual stimulation; during this period, the effect of word meaning gradually dissipates.

In the Auditory modality (Figure 1b), the different phases in the pupil response are not clearly separable, presumably because there is no visual stimulation to induce pupil constriction; instead, there is a uniform dilation that peaks after about 1,200 ms, and which is modulated by word meaning for a prolonged period (890 - 1850 ms).

To see how stable the effect was across participants, we determined, for every participant separately, the mean difference in pupil size between darkness- and brightness-related words in a 500 to 1,500 ms window (Figure 2). (This window deviates from the 1,000 to 2,000 ms window used in Mathôt et al. (2017); this deviation was a post-hoc decision based on the observed time course of the Visual experiment.) We then conducted a default Bayesian one-sample, one-tailed t-test to quantify the evidence for a difference in the predicted direction. For the Visual experiment, there was ‘substantial’ evidence for a difference (BF10 = 3.1). For the Auditory experiment, there was also ‘substantial’ evidence for a difference (BF10 = 8.0). (See e.g. Wetzels et al. (2011) for a qualitative interpretation of Bayes Factors.)

**Figure 2.**
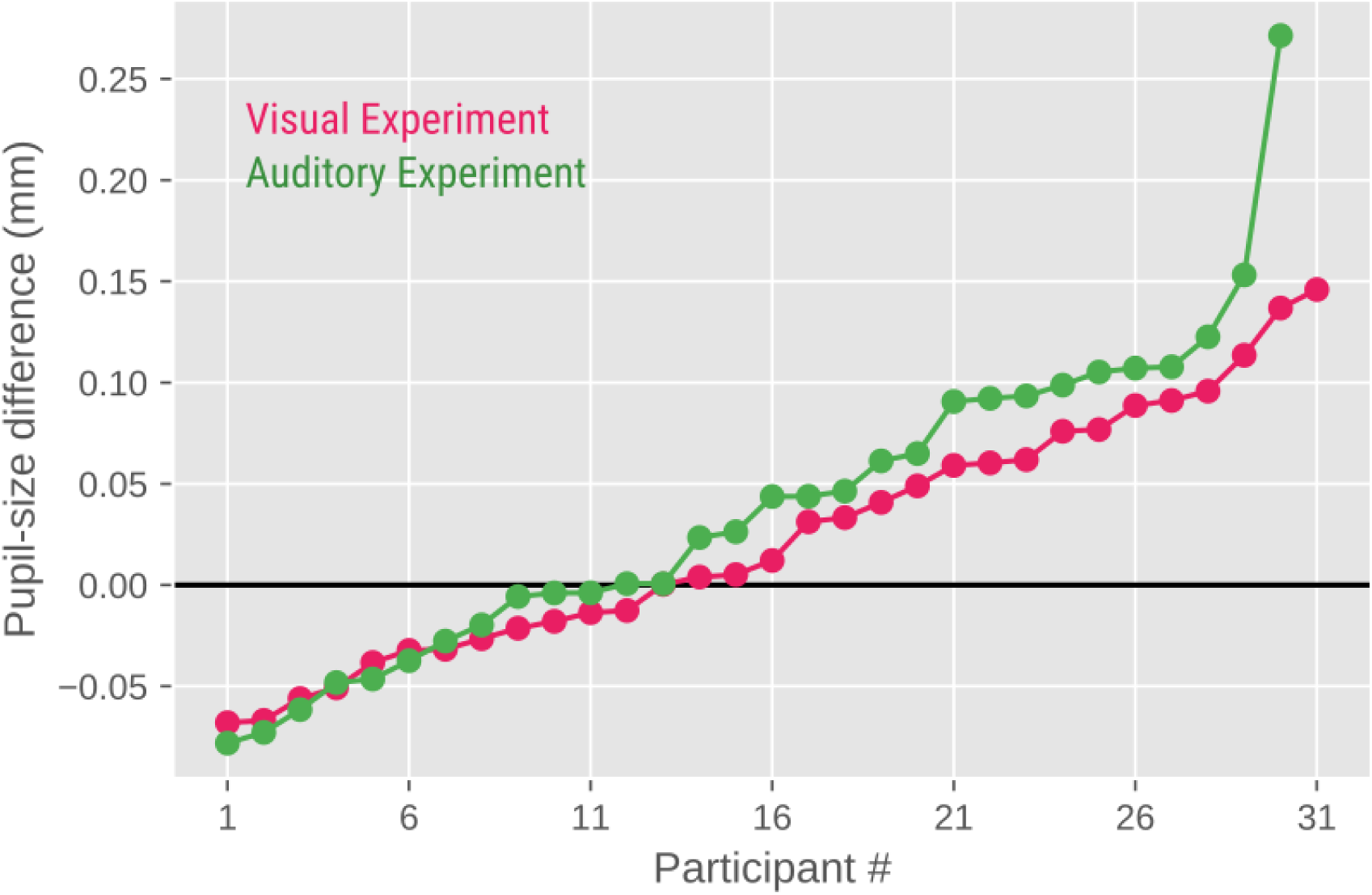
The mean difference in pupil size between darkness- and brightness-related words in the 500 - 1,500 ms window after word onset for the Visual (pink line) and Auditory (green line) experiment. Dots are individual participants, rank-ordered by effect size.

To inspect pupil responses to individual words, we determined, for every word separately, the mean pupil size in the 500 to 1,500 ms window. As shown in Figure 3, the difference between brightness- and darkness-related words is small compared to the variation within word categories; that is, semantic brightness explains only a small part of the variation in pupil size. However, there is a shift in the distribution, such that pupil size is overall larger in response to darkness-related words, as compared to brightness-related words. We then conducted a default Bayesian independent-samples, one-tailed t-test to quantify the evidence for a difference in the predicted direction. For the Visual experiment, there was only ‘anecdotal’ evidence for a difference (BF10 = 1.4). For the Auditory experiment, there was again ‘substantial’ evidence for a difference (BF10 = 4.3).

**Figure 3.**
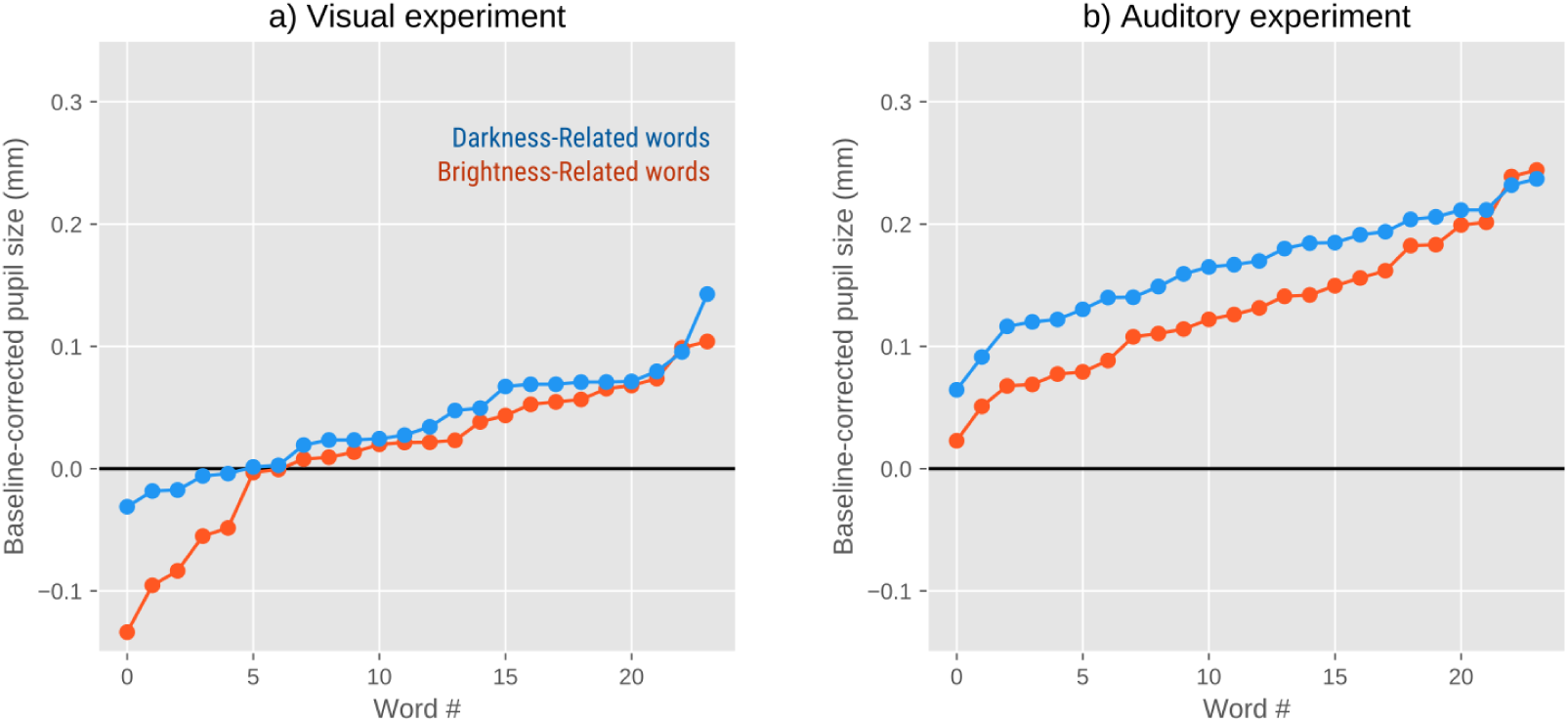
Mean pupil size for darkness-(blue lines) and brightness-related (orange lines) words in the 500 - 1,500 ms window after word onset, separately for the Visual (a) and Auditory (b) experiments. Dots are individual words, rank-ordered by pupil size.

Taken together, the results closely resemble those reported in Mathôt et al. (2017): the pupil is larger after seeing or hearing darkness-related words as compared to brightness-related words, and this effect builds up gradually and slowly. However, there are also notable differences. First, in the Visual experiment from Mathôt et al. (2017), the semantic-brightness effect peaked after about 1,800 ms, whereas in the current Visual experiment this effect peaks already after about 1,200 ms, and then dissipates. Second, the semantic-brightness effect was most pronounced in the Visual experiment from Mathôt et al. (2017), whereas it is most pronounced in the Auditory experiment from the current study.

### Normative ratings

A new group of 30 participants rated each of the words on three characteristics using a five-point scale: Semantic Brightness (very dark, dark, neither bright nor dark, bright, very bright), Concreteness (very abstract … very concrete), and Valence (very negative … very positive). We also determined the Emotional Intensity of each word by taking the absolute deviation from a neutral Valence, resulting in a three-point scale.

We first determined how Semantic Brightness correlated with the other word characteristics, using a Bayesian correlation analysis. The results roughly match those reported in Mathôt et al. (2017): a positive correlation between Semantic Brightness and Valence (*r* = .85, BF > 100), such that bright words were rated as more positive than dark words; a positive correlation between Semantic Brightness and Emotional Intensity (*r* = .37, BF = 4.7), such that bright words were rated as more intense than dark words; and no correlation between Semantic Brightness and Concreteness (*r* = -.06, BF = 0.20), such that bright and dark words were about equally concrete.

Our main analyses, described above, are based on a simple model that only considers Semantic Brightness as a predictor, and we relied on matching to eliminate the most likely confounds. However, we also conducted several supplementary analyses, described below, that included word characteristics as control predictors.

We controlled for Emotional Intensity and Concreteness by including these factors as control predictors in two separate linear mixed-effects models as described above (but originally run without control predictors). When including Emotional Intensity as control predictor, the effect of (rated) Semantic Brightness was still reliable for the Visual Experiment (850 - 1070 ms), but no longer for the Auditory Experiment (due to our criterion of having at least 200 contiguous milliseconds with *p* < .05). When including Concreteness as control predictor, the effect of (rated) Semantic Brightness was still reliable in both the Visual (850 - 1060 ms) and Auditory (750 - 2070 ms) experiments. Because Brightness and Valence are so highly correlated, we did not attempt to control for word Valence statistically. However, Mathôt et al. (2017) reported a control experiment in which word Valence was manipulated while Semantic Brightness was kept constant (which is much easier than the other way around), and found that word Valence does *not* affect pupil size.

In summary, when statistically controlling for Emotional Intensity and Concreteness, the effect of semantic brightness on pupil size largely remained reliable, with the exception of the Auditory experiment when controlling for Emotional Intensity. However, increased emotional intensity is associated with larger pupils (Bradley, Miccoli, Escrig, & Lang, 2008); therefore, the fact that bright words were rated as more emotionally intense than dark words should, if anything, have biased our results against our prediction.

## Discussion

We successfully replicated the finding, originally reported by Mathôt et al. (2017), that pupil size is affected by the semantic brightness of words; specifically, the pupil is slightly smaller after reading or hearing words that are associated with brightness, such as ‘sun’, as compared to words that are associated with darkness, such as ‘night’. The results of the current visual experiment were slightly different from those of the original experiment: In the current experiment we found only a brief effect (± 500 ms) of semantic brightness on pupil size, whereas the original experiment found a prolonged effect (± 2,000 ms). But overall the results of the current replication, conducted with Dutch words, resemble those of the original study, conducted with French words.

The present results confirm that the pupil light response (PLR) is an effective tool for the study of psycholinguistics. But what exactly does the effect of semantic brightness, or of higher level cognition in general, on the PLR reflect? To test this, Ebitz & Moore (2017) applied subthreshold microstimulation to the frontal eye fields (FEF), an area of the prefrontal cortex that is involved in the control of eye movements and visual attention (see also Wang & Munoz, 2018 for a similar study on the superior colliculus). When FEF cells are stimulated strongly (suprathreshold), an eye movement is triggered to a predictable location. But when FEF cells are stimulated weakly (subtreshold), a covert shift of attention is triggered to that same (stimulated) location, without a corresponding eye movement. Ebitz & Moore (2017) found that the PLR was enhanced when a bright stimulus was presented at a stimulated location, as compared to a control location. A plausible mechanistic explanation for this finding holds that the FEF feeds back to visual brain areas (Schall, Morel, King, & Bullier, 1995), where the representation of stimulated locations is enhanced; in turn, this might enhance the PLR to stimuli presented at stimulated locations. More generally, although many details are still unclear, it is likely that cognitive modulations of the PLR are a useful marker of activity in visual brain areas.

If the effect of semantic brightness on the PLR is robust, as the current replication suggests, and if it reflects activity in visual brain areas, then there are many psycholinguistic questions that could be addressed using this paradigm. For example, is a word processed differently depending on whether it is used metaphorically or not (cf. Sprenger, Levelt, & Kempen, 2006)? Is ‘dark’ processed differently in the sentences “I had no idea; I was totally in the dark” (metaphorical darkness) and “I turned off the light; I was totally in the dark” (real darkness)?

In summary, we successfully replicated the finding that the pupil is smaller after reading or hearing words that are associated with brightness, as compared to words that are associated with darkness. We have proposed that this effect is due to activity in visual brain areas, and can therefore be used as an indirect-yet-convenient marker for the recruitment of visual brain areas during language processing.

## Materials and availability

Data and experimental materials can found at https://osf.io/qd3vp/.

## References

Bárány, E. H., & Halldén, U. (1948). Phasic inhibition of the light reflex of the pupil during retinal rivalry. Journal of Neurophysiology, 11 (1), 25–30.

Binda, P., & Murray, S. O. (2014). Keeping a large-pupilled eye on high-level visual processing. Trends in Cognitive Sciences, 19(1), 1–3. http://doi.org/10.1016/j.tics.2014.11.002

Binda, P., Pereverzeva, M., & Murray, S. O. (2013a). Attention to bright surfaces enhances the pupillary light reflex. Journal of Neuroscience, 33(5), 2199–2204. http://doi.org/10.1523/jneurosci.3440-12.2013

Binda, P., Pereverzeva, M., & Murray, S. O. (2013b). Pupil constrictions to photographs of the sun. Journal of Vision, 13(6), e8. http://doi.org/10.1167/13.6.8

Bradley, M. M., Miccoli, L., Escrig, M. A., & Lang, P. J. (2008). The pupil as a measure of emotional arousal and autonomic activation. Psychophysiology, 45(4), 602–607. http://doi.org/10.1111/j.1469-8986.2008.00654.x

Campbell, F. W., & Gregory, A. H. (1960). Effect of size of pupil on visual acuity. Nature, 4743, 1121–1123. http://doi.org/10.1038/1871121c0

Dalmaijer, E., Mathôt, S., & Van der Stigchel, S. (2014). PyGaze: An open-source, cross-platform toolbox for minimal-effort programming of eyetracking experiments. Behavior Research Methods, 46(4), 913–921. http://doi.org/10.3758/s13428-013-0422-2

Ebitz, R. B., & Moore, T. (2017). Selective modulation of the pupil light reflex by prefrontal cortex microstimulation. Journal of Neuroscience, 2433–16. http://doi.org/10.1523/jneurosci.2433-16.2017

Ebitz, R. B., Pearson, J. M., & Platt, M. L. (2014). Pupil size and social vigilance in rhesus macaques. Frontiers in Decision Neuroscience, 8, 100. http://doi.org/10.3389/fnins.2014.00100

Ellis, C. J. (1981). The pupillary light reflex in normal subjects. British Journal of Ophthalmology, 65(11), 754–759. http://doi.org/10.1136/bjo.65.11.754

Fahle, M. W., Stemmler, T., & Spang, K. M. (2011). How much of the “unconscious” is just pre-threshold? Frontiers in Human Neuroscience, 5. http://doi.org/10.3389/fnhum.2011.00120

Glenberg, A. M., & Gallese, V. (2012). Action-based language: A theory of language acquisition, comprehension, and production. Cortex, 48(7), 905–922. http://doi.org/10.1016/j.cortex.2011.04.010

Glenberg, A. M., & Kaschak, M. P. (2002). Grounding language in action. Psychonomic Bulletin & Review, 9(3), 558–565. http://doi.org/10.3758/bf03196313

Harms, H. (1937). Ort und Wesen der Bildhemmung bei Schielenden. Graefe’s Archive for Clinical and Experimental Ophthalmology, 138(1), 149–210. http://doi.org/10.1007/bf01854538

Husta, C., Dalmaijer, E., Belopolsky, A., & Mathot, S. (2018). The pupillary light response reflects visual working memory content. bioRxiv, 477562. http://doi.org/10.1101/477562

Keuleers, E., Diependaele, K., & Brysbaert, M. (2010). Practice Effects in Large-Scale Visual Word Recognition Studies: A Lexical Decision Study on 14,000 Dutch Mono- and Disyllabic Words and Nonwords. Frontiers in Psychology, 1. http://doi.org/10.3389/fpsyg.2010.00174

Krause, F., & Lindemann, O. (2014). Expyriment: A Python library for cognitive and neuroscientific experiments. Behavior Research Methods, 46(2), 416–428. http://doi.org/10.3758/s13428-013-0390-6

Laeng, B., & Endestad, T. (2012). Bright illusions reduce the eye’s pupil. Proceedings of the National Academy of Sciences, 109(6), 2162–2167. http://doi.org/10.1073/pnas.1118298109

Laeng, B., & Sulutvedt, U. (2014). The eye pupil adjusts to imaginary light. Psychological Science, 25(1), 188–197. http://doi.org/10.1177/0956797613503556

Lombardo, M., & Lombardo, G. (2010). Wave aberration of human eyes and new descriptors of image optical quality and visual performance. Journal of Cataract & Refractive Surgery, 36(2), 313–331. http://doi.org/10.1016/j.jcrs.2009.09.026

Lowe, S. W., & Ogle, K. N. (1966). Dynamics of the pupil during binocular rivalry. Archives of Ophthalmology, 75(3), 395. http://doi.org/10.1001/archopht.1966.00970050397017

Mahon, B. Z., & Caramazza, A. (2008). A critical look at the embodied cognition hypothesis and a new proposal for grounding conceptual content. Journal of Physiology-Paris, 102(1–3), 59–70. http://doi.org/10.1016/j.jphysparis.2008.03.004

Mathôt, S. (2013). A Simple Way to Reconstruct Pupil Size During Eye Blinks. Retrieved from http://dx.doi.org/10.6084/m9.figshare.688001

Mathôt, S. (2018). Pupillometry: Psychology, physiology, and function. Journal of Cognition, 1(1), 1–16. http://doi.org/10.5334/joc.18

Mathôt, S., & Van der Stigchel, S. (2015). New light on the mind’s eye: The pupillary light response as active vision. Current Directions in Psychological Science, 24(5), 374–378. http://doi.org/10.1177/0963721415593725

Mathôt, S., Dalmaijer, E., Grainger, J., & Van der Stigchel, S. (2014). The pupillary light response reflects exogenous attention and inhibition of return. Journal of Vision, 14(14), 7. http://doi.org/10.1167/14.14.7

Mathôt, S., Fabius, J., Heusden, E. V., & Stigchel, S. V. D. (2018). Safe and sensible preprocessing and baseline correction of pupil-size data. Behavior Research Methods, 1–13. http://doi.org/10.3758/s13428-017-1007-2

Mathôt, S., Grainger, J., & Strijkers, K. (2017). Pupillary responses to words that convey a sense of brightness or darkness. Psychological Science, 28(8), 1116–1124. http://doi.org/10.1177/0956797617702699

Mathôt, S., Schreij, D., & Theeuwes, J. (2012). OpenSesame: An open-source, graphical experiment builder for the social sciences. Behavior Research Methods, 44(2), 314–324. http://doi.org/10.3758/s13428-011-0168-7

Mathôt, S., van der Linden, L., Grainger, J., & Vitu, F. (2013). The pupillary response to light reflects the focus of covert visual attention. PLoS ONE, 8(10), e78168. http://doi.org/10.1371/journal.pone.0078168

Mathôt, S., van der Linden, L., Grainger, J., & Vitu, F. (2015). The pupillary light response reflects eye-movement preparation. Journal of Experimental Psychology: Human Perception and Performance, 41(1), 28–35. http://doi.org/10.1037/a0038653

Naber, M., & Nakayama, K. (2013). Pupil responses to high-level image content. Journal of Vision, 13(6), e7. http://doi.org/10.1167/13.6.7

Naber, M., Alvarez, G. A., & Nakayama, K. (2013). Tracking the allocation of attention using human pupillary oscillations. Frontiers in Psychology, 4(919), 1–12. http://doi.org/10.3389/fpsyg.2013.00919

Naber, M., Frassle, S., & Einhauser, W. (2011). Perceptual rivalry: Reflexes reveal the gradual nature of visual awareness. PloS ONE, 6(6), e20910. http://doi.org/10.1371/journal.pone.0020910

Papesh, M. H. (2015). Just out of reach: On the reliability of the action-sentence compatibility effect. Journal of Experimental Psychology. General, 144(6), e116–141. http://doi.org/10.1037/xge0000125

Pulvermüller, F. (2013). How neurons make meaning: brain mechanisms for embodied and abstract-symbolic semantics. Trends in Cognitive Sciences, 17(9), 458–470. http://doi.org/10.1016/j.tics.2013.06.004

Schall, J. D., Morel, A., King, D. J., & Bullier, J. (1995). Topography of visual cortex connections with frontal eye field in macaque: convergence and segregation of processing streams. Journal of Neuroscience, 15(6), 4464–4487. http://doi.org/10.1523/jneurosci.15-06-04464.1995

Schwanenflugel, P. J. (2013). Why are abstract concepts hard to understand? In The Psychology of Word Meanings (pp. 223–250). Routledge.

Sperandio, I., Bond, N., & Binda, P. (2018). Pupil size as a gateway Into conscious interpretation of brightness. Frontiers in Neurology, 9. http://doi.org/10.3389/fneur.2018.01070

Sprenger, S. A., Levelt, W. J., & Kempen, G. (2006). Lexical access during the production of idiomatic phrases. Journal of Memory and Language, 54(2), 161–184. http://doi.org/10.1016/j.jml.2005.11.001

Wang, C., & Munoz, D. P. (2015). A circuit for pupil orienting responses: implications for cognitive modulation of pupil size. Current Opinion in Neurobiology, 33, 134–140. http://doi.org/10.1016/j.conb.2015.03.018

Wang, C., & Munoz, D. P. (2018). Neural basis of location-specific pupil luminance modulation. Proceedings of the National Academy of Sciences, 115(41), 10446–10451. http://doi.org/10.1073/pnas.1809668115

Watson, A. B. (2013). A formula for the mean human optical modulation transfer function as a function of pupil size. Journal of Vision, 13(6), 18. http://doi.org/10.1167/13.6.18

Wetzels, R., Matzke, D., Lee, M. D., Rouder, J. N., Iverson, G. J., & Wagenmakers, E. J. (2011). Statistical evidence in experimental psychology. Perspectives on Psychological Science, 6(3), 291–298. http://doi.org/10.1177/1745691611406923

